# Heritability of the *Symbiodinium* community in vertically- and horizontally-transmitting broadcast spawning corals

**DOI:** 10.1101/100453

**Authors:** Kate Quigley, Bette Willis, Line Bay

## Abstract

The dinoflagellate-coral partnership influences the coral holobiont’s tolerance to thermal stress and bleaching. However, the comparative roles of host genetic versus environmental factors in determining the composition of this symbiosis are largely unknown. Here we quantify the heritability of the initial *Symbiodinium* communities for two broadcast-spawning corals with different symbiont transmission modes: *Acropora tenuis* has environmental acquisition, whereas *Montipora digitata* has maternal transmission. Using high throughput sequencing of the ITS-2 region to characterize communities in parents, juveniles and eggs, we describe previously undocumented *Symbiodinium* diversity and dynamics in both corals. After one month of uptake in the field, *Symbiodinium* communities associated with *A. tenuis* juveniles were dominated by A3, C1, D1, A-type CCMP828, and D1a in proportional abundances conserved between experiments in two years. *M. digitata* eggs were predominantly characterized by C15, D1, and A3. In contrast to current paradigms, host genetic influences accounted for a surprising 29% of phenotypic variation in *Symbiodinium* communities in the horizontally-transmitting *A. tenuis*, but only 62% in the vertically-transmitting *M. digitata*. Our results reveal hitherto unknown flexibility in the acquisition of *Symbiodinium* communities and substantial heritability in both species, providing material for selection to produce partnerships that are locally adapted to changing environmental conditions.

## Introduction

Coral bleaching, defined as either the loss of *Symbiodinium* cells from coral tissues or reduction in symbiont photosynthetic pigments, represents a threat to coral reefs world-wide as it increases in both frequency and magnitude ^1–4^ If coral reefs are to persist under climate change, corals must either disperse to new unaffected habitats, acclimate through phenotypic plasticity, and/or adapt through evolutionary mechanisms ^5^. However, the extent to which thermal tolerance can increase, either through changes to the host genome or *Symbiodinium* community hosted, or by direct selection on the symbionts themselves, is currently unclear.

Bleaching sensitivity is variable within and among species ^6^, but comparative roles of host genetics versus symbiont communities to this variation remain unclear ^7,8^. The *Symbiodinium* community hosted by corals has long been recognized as the primary factor. determining bleaching susceptibility ^8,9^. However, host influences are also evident ^10–12^ and may play an equally important role in determining bleaching susceptibility. Endosymbiotic communities could influence host adaptation to changing climates through increased host niche expansion ^13,14^, but a major impediment to understanding the capacity of corals to adapt to a changing climate is lack of knowledge about the extent to which *Symbiodinium* communities associated with corals are inherited and hence subject to selection.

There are nine recognized *Symbiodinium* clades ^15^ that encompass substantial sequence and functional variation at the intra-clade (type) level (reviewed in ^16^). Deep sequencing technologies currently available can detect type level diversity even at low abundances ^17^ and are now being applied to understand adult coral-*Symbiodinium* diversity ^18–20^, but have not yet been applied to the early life-history stages of corals. Therefore, there are gaps in our basic knowledge of the composition of *Symbiodinium* communities at lower, functionally relevant taxonomic levels, particularly community members at background abundances, and in the eggs and juveniles of corals.

Natural variation in the composition of coral-associated *Symbiodinium* communities exists among coral populations and species ^16,21^, with certain communities offering greater bleaching resistance compared to others ^22,23^. It is not yet known what enhances or constrains the capacity of corals to harbour stress-tolerant *Symbiodinium* types and whether changes in *Symbiodinium* communities in response to environmental stressors are stochastic or deterministic ^24^. Given the importance of *Symbiodinium* communities for bleaching susceptibility and mortality of the coral holobiont ^25,26^, quantifying the proportional contributions of genetic and environmental factors to community formation, regulation and stress tolerance is important for understanding coral health. If the *Symbiodinium* community is heritable, changes to these communities may bring about adaptation of the holobiont as a whole. Under this scenario, *Symbiodinium* community shifts are equivalent to changes in host allele frequencies, thus opening up new avenues for natural and artificial selection, assisted evolution and microbiome engineering ^24,27^.

*Symbiodinium* communities associated with scleractinian corals are either acquired from the environment (horizontal transfer) or passed maternally from adults to eggs or larvae (vertical transfer). Approximately 85% of scleractinian coral species broadcast spawn eggs and sperm into the environment, and of these, ~80% acquire symbionts horizontally; the remaining ~20% acquire them vertically ^28^. Vertically-transmitted symbiont communities are predominantly found in brooding corals with internal fertilization ^28^ and are theorized to be of lower diversity and higher fidelity ^16^^16^. Conversely, horizontal transmission has generally been assumed to result in weaker fidelity that can be increased through the development of strong genotype associations between hosts and their symbiont community ^29^. Studies specifically quantifying the genetic component governing *Symbiodinium* communities established in offspring of both horizontal and vertical transmitters are needed to elucidate the potential for adaptation through symbiont community changes.

Heritability describes the genetic components of variability in a trait using analysis of co-variance among individuals with different relatedness ^30^. The ratio of additive genetic variance to phenotypic variance (V_A_/V_P_) is defined as narrow-sense heritability (h^2^) ^31^. The degree of heritability of a trait ranges from 0 - 1, and describes the influence of parental genetics on the variability of that trait ^31^. Therefore, the degree to which traits might change from one generation to the next can be predicted from measures of heritability, where the predicted change in offspring phenotype is proportional to h^2^ (i.e., the breeder’s equation) ^32^. It is particularly important to determine the genetic contribution to understand the potential for adaptation and to predict the strength of response to selection (i.e, the ‘evolvability’ of a trait) ^5,33,34^.

To quantify the potential for selection of endosymbiotic *Symbiodinium* communities associated with broadcast spawning corals in response to changes in environmental conditions (i.e., climate change-induced), we characterized symbiont communities associated with adults and juveniles of the horizontal transmitter *Acropora tenuis* and with adults and eggs of the vertical transmitter *Montipora digitata* using high-throughput sequencing. Using a community diversity metric, we derived the narrow-sense heritability (h^2^) of these communities and identified new and unique *Symbiodinium* types recovered from juveniles and eggs compared to their parental colonies. Finally, we described previously unknown *Symbiodinium* community dynamics in the early life-history stages of these two common coral species.

## Results

### *Symbiodinium* communities associated with *Acropora tenuis*

After one month in the field, there were similarities at the clade level between *Symbiodinium* communities associated with the 2012 and 2013 families of *A. tenuis* juveniles, with 54 OTUs (17.1%) shared between the two years, including similar proportions of OTUs retrieved across the clades in each year (Fig. 1, Supplementary Table S4). In both years, the majority of OTUs were recovered from three clades (A, C, and D) and the number of OTUs from each of these clades was similar between years (Supplementary Table S4). The greatest diversity of OTUs found in juveniles from both years belonged to C1, A3 and “uncultured” types (see methods for definitions of OTUs), and a diversity of different OTUs within types A13, A-type CCMP828, D1 and D1a were also present (Supplementary Fig. S1). The predominant patterns characterising *Symbiodinium* communities associated with the 2012 and 2013 families were the high abundance of *Symbiodinium* types A3, C1, D1, and CCMP828, and the comparatively lower abundance of D1a (Fig. 2). However, substantial variation in *Symbiodinium* diversity and abundance existed among juveniles within the same family, as well as among families of juveniles (Supplementary Results, Fig. 2). For example, juvenile families differed in their average OTU diversity and abundance, as well as their taxonomic composition (additional description in Supplementary Results, Fig. 2, Supplementary Table S5), where particular families contained juveniles of particularly high diversity (families F14 and F18).

**Figure. 1.**
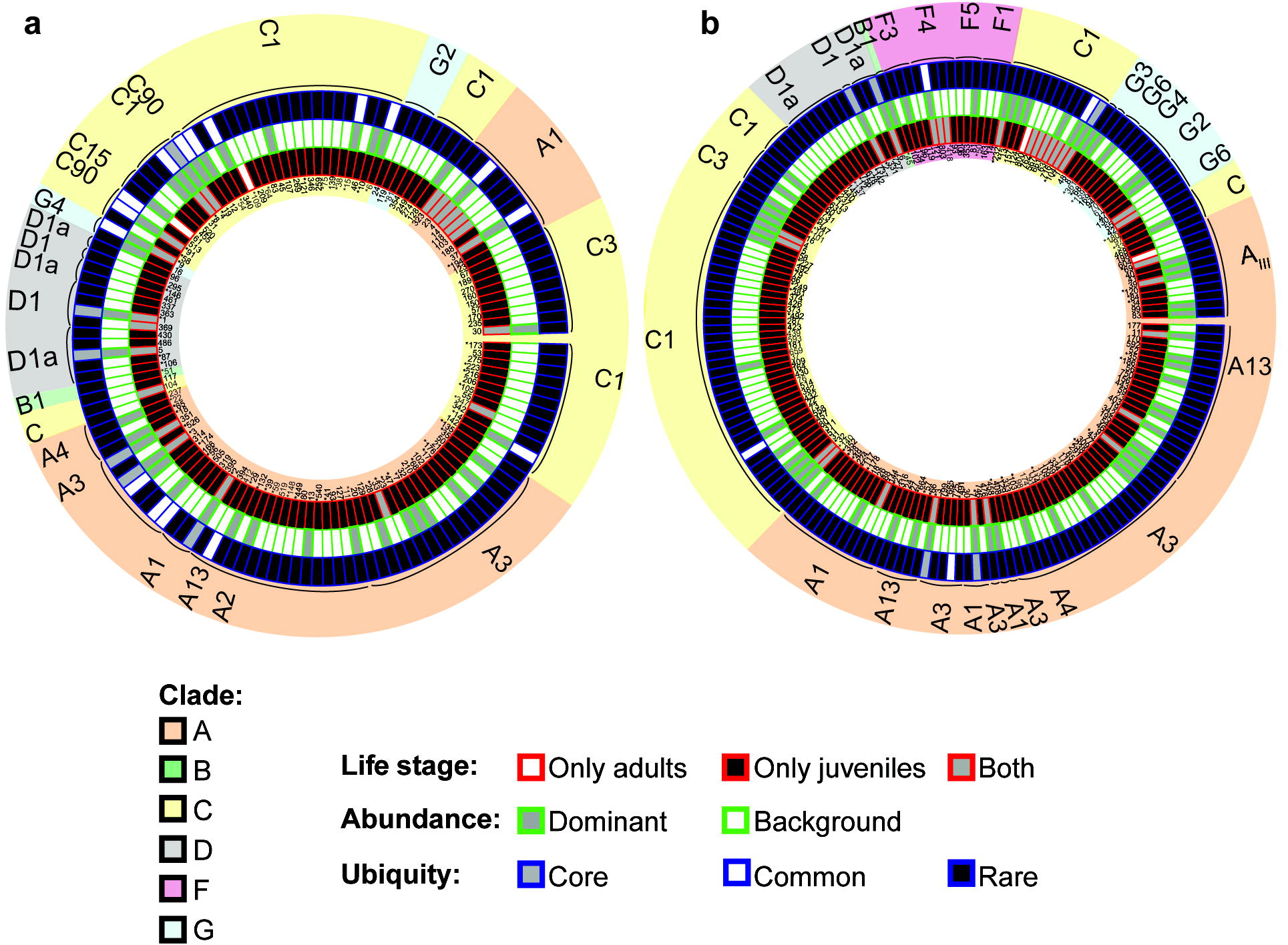
Circular trait plots of 261 *Symbiodinium* ITS-2 OTUs retrieved from *Acropora tenuis* juveniles and adults in 2012 (a) and 2013 (b). Plots include only those OTUs that were retrieved from three or more samples (134/422 OTUs in 2012 and 181/568 OTUs in 2013). Concentric circles from innermost to the outermost position represent OTUs present: 1) life stage, 2) normalized abundance (principal: > 0.01%, background < 0.01%), and 3) ubiquity (core: >75% of samples, common: 25-75%, rare: < 25%). OTU identity with an asterisk indicates it was retrieved in both years. Semi-transparent backgrounds represent clade designations of individual OTUs. See Supplementary Table S8 for full taxonomic information.

**Figure. 2.**
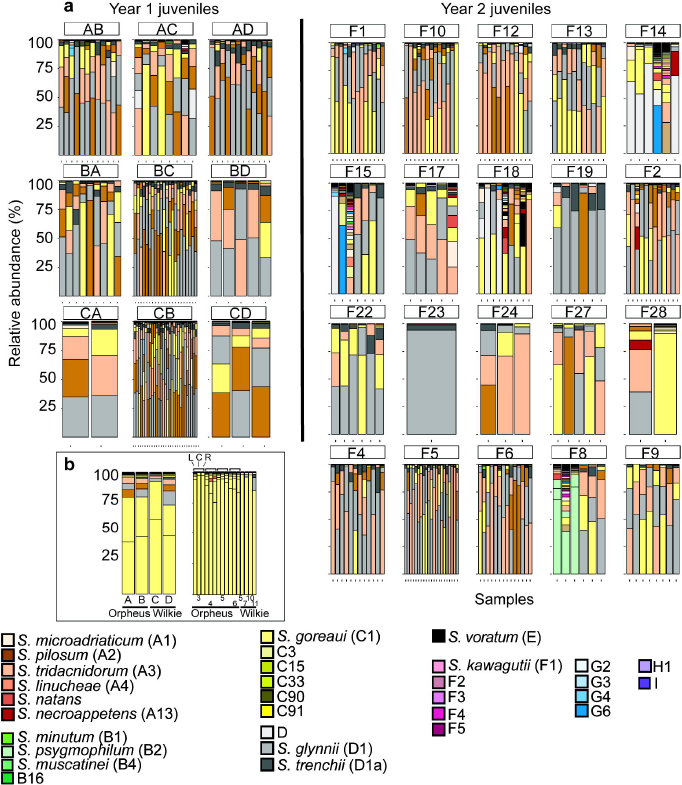
Barplots of variance-normalized abundances of *Symbiodinium* diversity associated with (a) juveniles and (b) adults of *Acropora tenuis* used in 2012 (Year 1) and 2013 (Year 2) crosses. Colours represent different *Symbiodinium* types. Origins of parent colonies are Orpheus and Wilkie reefs. *A. tenuis* adult colonies from Orpheus used for 2013 crosses included samples that were sequenced that represent the left side of the colony (L), center of the colony (C), and right side of the colony (R) to examine intra-colony *Symbiodinium* diversity.

Juveniles from both years harboured more unique OTUs than adults (juveniles vs. adults: 111 vs. 2 (2012), 151 vs. 2 (2013)), with comparatively few OTUs shared between life stages (21 shared in 2012 (out of 422 OTUs); 28 shared in 2013 (out of 568 OTUs)) (Fig. 1). Furthermore, the majority of OTUs in both years were at background abundances (Fig. 1). The majority of OTUs were also rare (112 - 172 OTUs found in less than 25% of samples in 2012 and 2013), whilst 4 - 16 OTUs were common (25 -75% of samples) and 5 - 6 OTUs were core members (two A3 types, CCMP828, C1, D1, D1a were present in greater than 75% of samples) (Fig. 1).

### *Symbiodinium* communities associated with *Montipora digitata*

101 OTUs were found in *M. digitata* eggs and adults, with 7 (±0.9 SE) OTUs per egg and 5.3 (±0.9 SE) OTUs per adult, on average. The highest diversities of OTUs were retrieved from clades A (73 OTUs) and C (18 OTUs), whereas D had three OTUs represented (Fig. 3). 99.1% of the total cleaned reads belonged to C15 (OTU1), with this type making up 98.8 % (±0.5 SE) and 99 % (±0.1 SE) of all reads retrieved from dams and eggs, respectively. The next most abundant OTUs were C1, D1, and A3 (Fig. 4). Adults could generally be distinguished from eggs by the unique presence of A2, A3, particular C1 and A3 variants (C1_8, HA3-5), G3 (Fig. 3), and a greater proportional abundance of an A type symbiont (OTU4) in dams 29, 32, 7, 8 and 9 (Fig. 4). Of these unique adult OTUs, none were found in more than two adult colonies. Eighty-two OTUs were found in eggs but not adults and 43 of these were found in three or more eggs, and a majority were “uncultured” types at background levels from the eggs of dam 29 (Fig. 3). Both inter and intra family variation in background *Symbiodinium* OTU composition and abundance were detected within eggs as well (further description in Supplementary Results, Supplementary Fig. S2, Supplementary Table S6).

**Figure. 3.**
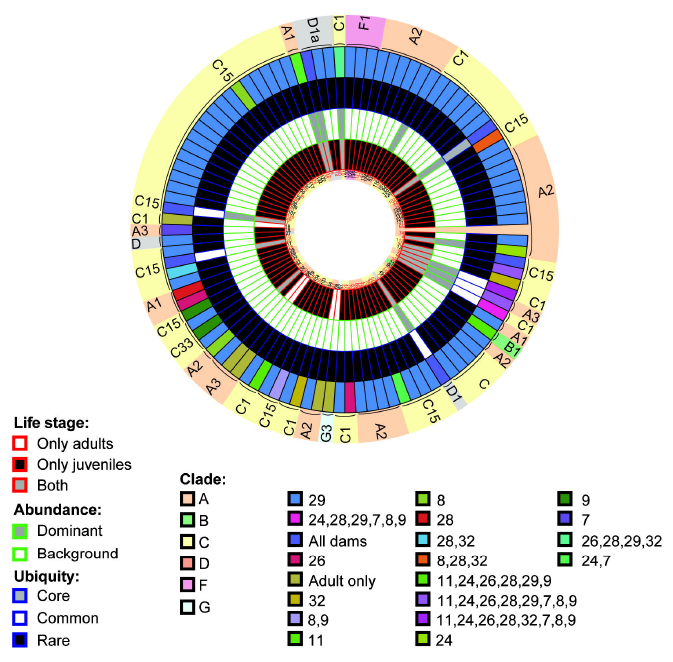
Circular trait plots of 101 *Symbiodinium* ITS-2 OTUs retrieved from *Montipora digitata* eggs and adults. Concentric circles from innermost to the outermost position represent OTUs present: 1) life-stage, 2) normalized abundance (principal: > 0.01%, background < 0.01%), 3) ubiquity (core: >75% of samples, common: 25-75%, rare: < 25%), and 4) dam identity. Semi-transparent backgrounds represent clade designations of individual OTUs. Red text indicates OTUs that were found in three or more eggs or adults. See Supplementary Table S8 for full taxonomic information.

**Figure. 4.**
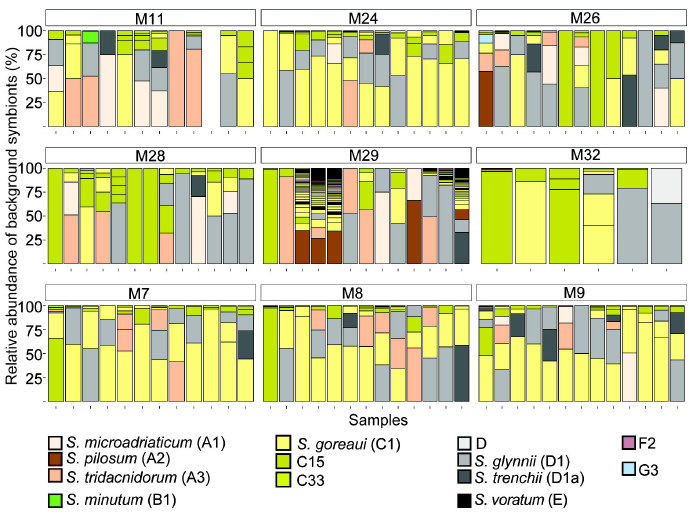
Barplot of variance-normalized abundances of only the background *Symbiodinium* diversity associated with dams and eggs of *Montipora digitata.* Colours represent different *Symbiodinium* types. The dominant type, C15, was excluded for clarity. The first bar in each group is the spawning dam and the following bars represent her eggs. The tenth egg sample from dam 11 (M11) was made up of 100% C15, and was therefore not shown.

### Narrow-sense heritability of *Symbiodinium* community in *A. tenuis* juveniles and *M. digitata* eggs

Bayesian linear mixed models, and specifically, the animal model, were used to estimate relatedness-based heritability as they are robust to unbalanced designs. Furthermore, the animal model utilizes all levels of relatedness between individuals in a given dataset, and not just parent-offspring comparisons ^35^. The Bayesian narrow-sense heritability estimate (h^2^) of the initial *Symbiodinium* community in *A. tenuis* juveniles was 0.29, with a 95% Bayesian credibility interval for the additive genetic component of 0.06-0.86. The mean heritability was 0.36 (±0.21 SD) (Fig. 5). The high density of estimates between 0.2 - 0.4 within the posterior distribution of h^2^ suggests high statistical support around 0.29, despite the credibility interval being very large. The maternal transfer of *Symbiodinium* in the broadcast spawning coral *M. digitata* had a narrow-sense heritability estimate of 0.62 (0.27-0.86 95% Bayesian credibility interval), with a mean heritability of 0.57 (±0.16 SD) (Fig. 5). We did not detect an effect of maternal environment on similarities in *Symbiodinium* diversity among eggs or among juveniles. Models that included maternal effects arising from eggs developing in a shared environment (maternal environmental effects for both *A. tenuis* and *M. digitata*) were not significantly better than those that did not include maternal effects (DIC no effects < DIC maternal environmental effects included).

**Figure. 5.**
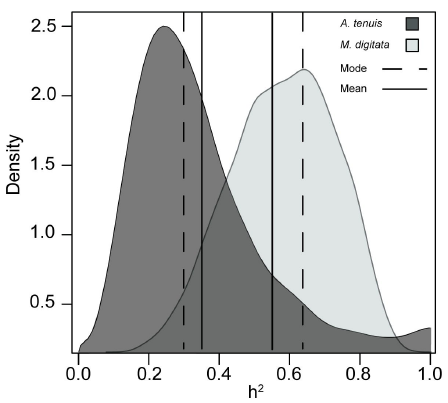
Posterior distributions of the heritability estimates for *A. tenuis* (dark grey) and *M. digitata* (light grey) generated from Bayesian MCMCglmm models. Dashed and full lines correspond to distribution modes and means, respectively.

Mid-parent regression estimates for the 29 *A. tenuis* families from 2012 and 2013 indicated that trait-based h^2^ of the *Symbiodinium* community was 0.3 (Supplementary Fig.S3). Parent-offspring regression of the 99 *M. digitata* eggs genotyped from nine dams resulted in a heritability estimate of 0.16 (slope= 0.078 x 2 as a single parent) (Supplementary Fig. S4). Therefore, 30% and 16% of the measured variation in the *Symbiodinium* community in *A. tenuis* and *M. digitata,* respectively, was due to genetic differences among offspring.

### Impact of intragenomic variation on heritability analysis

Simulating intragenomic variants in the *M. digitata* dataset yielded five intragenomic variant groups from clade A (IGV1_A: OTU_65/74/113/123/121; IGV2_A: 29/23/133; IGV3_A: 68/32; IGV4_A:61/70; IGV5_A: 56/75), and one from clade C (IGV6_C: 128/42). OTUs from clade D were not highly similar and correlation coefficients for the three clade F OTUs had relatively low correlation coefficients (0.3-0.6). The diversity metric and Bayesian MCMC heritability was re-calculated with these 16 OTUs collapsed into their respective six intragenomic variants. The resulting h^2^ estimate was slightly higher (0.5754±0.157 compared to the original estimate of 0.5722±0.157).

## Discussion

Substantial heritability of the initial *Symbiodinium* community in early life history stages of both vertically and horizontally-transmitting corals highlights the important role of host genetics in governing the composition of symbiont communities within their tissues. Surprisingly, mean Bayesian heritability estimates for initial *Symbiodinium* communities associated with juveniles of *Acropora tenuis* were moderate (0.29), but higher than expected given low levels of fidelity assumed for species with environmentally-acquired symbionts. Conversely, heritability estimates associated with eggs of *Montipora digitata* were high (0.62), but lower than expected given the high levels of fidelity expected for vertically transmitted symbionts. Given that heritability is a quantifiable measure of the influence of genes compared to environmental factors in shaping phenotypes, both non-zero heritability estimates confirm that genes do influence the structuring of *Symbiodinium* communities in these two coral species. Although our results differ from expectations of fidelity and heritability based on current transmission paradigms in corals, they are consistent with studies that have demonstrated the role of host genetics in governing the composition of symbiotic bacterial communities in mammals, insects and other cnidarians ^36–39^, as well as the abundance of bacteria in insects ^40^ and humans ^41^. Furthermore, these estimates are consistent with the characteristic hallmarks of host-controlled symbiont regulation. For example, *Symbiodinium* cells are enveloped in a host-derived symbiosome, with only a few (2-8) symbiont cells per host membrane ^42^. This suggests that the coral host may regulate *Symbiodinium* on an almost individual cell basis, facilitating overall population regulation ^40^ and potentially community composition within the holobiont. Thus, it is likely advantageous for the host’s molecular architecture governing the *Symbiodinium* community to be passed from one generation to the next. Importantly, the partial genetic regulation of *Symbiodinium* communities found here suggests that there is potential for the symbioses to evolve and adapt, and therefore to potentially develop ‘optimal’ symbiont-host partnerships under changing environmental conditions.

Our results provide the first in-depth picture of the complexity of the *Symbiodinium* community in *A. tenuis* juveniles during the initial month of uptake. No juveniles exclusively hosted a single clade or type, a result corroborated by lab and other field-based experimental studies ^43–48^. Moreover, although the diversity measured here was much greater than values reported in previous studies, we found temporal stability in cladal diversity and abundances between the two years. It is possible that the temporal stability detected at the clade level within juveniles was in part due to the stability of locally available symbionts, either from the sediments or from the continual seeding of symbionts into local environments by resident symbiont-bearing cnidarians ^49^. Such environmental variance is partitioned in the MCMC animal model (along with genetic effects due to relatedness) and hence accounted for in heritability estimates. Therefore, stability in the availability of environmental *Symbiodinium* and its subsequent impact on temporal stability of coral-associated *Symbiodinium* communities would be accounted for in heritability estimates. The unexpectedly high fidelity of the symbiont community, in conjunction with our heritability estimates, suggest strong host genetic – symbiont community associations, a result also implicated in studies comparing symbiosis fidelity across phylogenetic associations in *Hydra,* wasps, and primates ^29^. Further work is needed to document *Symbiodinium* diversity in juveniles of broadcast spawning corals, as well as to elucidate molecular mechanisms regulating the establishment of this symbiosis.

Our conclusion of active host regulation based on heritability estimates, coupled with temporal stability in the relative proportions and numbers of OTUs within clades at principal and background levels between years, suggest that genetic regulation governing *Symbiodinium* communities extends to clades found at very low abundance. The roles of many background *Symbiodinium* types remain unclear and may be minor compared to principal types like A3, C1, and D1 when corals are healthy. However, *Symbiodinium* at background abundances can be important for coral health under sub-optimal environmental conditions. For example, fine scale dynamics of *Symbiodinium* communities (i.e., changes in relative abundance and/or diversity of only a fraction of types) impact host bleaching susceptibility, recovery and physiology ^23,26,50,51^. Growing evidence suggests that background types are important in several *Symbiodinium-coral* symbioses during recovery from stress (i.e. *Acropora millepora* and D-types ^23^, *Agaricia* spp., *M. annularis, M. cavernosa-D1a* ^50,52^, *Pocillopora damicornis, Stylophora pistilata*-C_I:53 ^19^), but may not be relevent for all (i.e. *Acropora japonica-* and *S. voratum* ^53^). A strong functional role of background *Symbiodinium* types would not be surprising given the functional importance of background bacterial lineages recently described for corals ^54,55^, but remains to be conclusively established for many *coral-Symbiodinium* associations.

The heritability signal derived from Bayesian models found for *Symbiodinium* communities associated with eggs of the vertically-transmitting coral *M. digitata* was predictably strong (62%) given that the dominant C15 OTU was harboured in adults and eggs at very high abundances. However, fidelity was less than expected given that eggs acquire *Symbiodinium* communities in the maternal environment. This lower than expected heritability signal is mirrored when the likenesses between dams and eggs are compared. For example, despite *Symbiodinium* C15 dominating symbiont communities in both eggs and dams, maternal transfer lacked precision in one dam in particular (dam 29), whose eggs had highly variable *Symbiodinium* communities that included “uncultured” OTUs, similar to previous reports for another species in this genus ^56^. It should be noted that, on average, *M. digitata* dams transmitted C15 so that it comprised 99% of the *Symbiodinium* community in all eggs, suggesting that a larger heritability estimate might have been expected. The posterior distribution of the heritability estimates also suggests that the value could resolve to be larger with increased sampling (up to 0.85). However, the lower than expected heritability reflects the fact that all *Symbiodinium* in eggs were considered, not just the OTU in greatest abundance. Therefore, the presence of other symbionts (although in low abundance and number) lowered the heritability estimate. The incorporation of all *Symbiodinium,* and not just the numerically dominant one, in heritability calculations is ecologically relevant, given the important role that low abundance microbes have in coral physiology and stress tolerance (clade D *Symbiodinium* ^57,58^, and bacteria ^54,55^).

There are many precedents for inexact maternal transfer of symbiont communities, and studies on insects show that vertical transmission is rarely perfect ^59^ due to symbiont competition within hosts ^60^. Such imprecision in maternal transfer is a product of fitness costs associated with the maintenance of superinfections (stable coexistence of multiple symbionts) and can be overcome if selection for coexistence is greater than costs associated with their maintenance ^60^. Superinfections may provide a diversity of beneficial symbiont traits. For example, different symbionts provide different nutrients to host insects ^61^. For *M. digitata,* imprecision may represent a bet-hedging strategy to maximise the likelihood that some offspring will survive when eggs are dispersed and encounter environments that are different to their parents. Although some of these background OTUs may represent random contaminants (i.e. symbionts attached to the outside of eggs), a majority of OTUs were found in three or more independent egg samples, suggesting that they indeed represent either relevant symbiont candidates or intragenomic variants retrieved from relevant symbiont candidates. However, it is unlikely that these OTUs are intragenomic variants given the clustering method and clustering identity threshold used in this study (Materials and Methods: Sequencing of *Symbiodinium* ITS-2 in egg, juvenile and adult coral samples). Although many of these background OTUs existed predominantly at less than 1% abundance in adults and eggs, it is feasible that these OTUs may grow in abundance to become dominant members of the community if environmental conditions change ^50^, as was found for C.28 and C_I:53 in *P. damicornis* ^19^. This variation highlights potential flexibility in the *M. digitata Symbiodinium* symbiosis, which may enable the host to vary its symbiotic partnerships in response to environmental change by benefitting from new host-symbiont combinations.

Surprisingly, much of the diversity found in *M. digitata* eggs was not present in parent colonies, similar to results reported for larvae of the brooding, vertically-transmitting coral *Seriatopora hystrix* (Quigley et al. *in-review*) and observed here between *A. tenuis* juveniles and adults (this study). Our results suggest that eggs acquire symbionts from sources external to the maternal transmission process. Mixed systems involving both vertical and horizontal transmission are known (e.g. bacteria in clams; reviewed in ^29^) and have recently been demonstrated in brooding corals (Quigley et al. *in-review).* Given that the cellular machinery needed for recognition of appropriate *Symbiodinium* types ^42^ would not be developed in egg cytoplasm, where *Symbiodinium* are present pre-fertilization ^62^, eggs exposed to transient symbionts in the dam’s gastrovascular cavity or by parasitic *Symbiodinium-containing* vectors (e.g. ciliates ^63^ and parasites ^60^) may retain these communities until recognition systems of eggs, larvae or juveniles mature. Interestingly, one type (OTU111) found in three eggs from dam 29 was identified as a free-living A type recovered from Japanese marine sediments (EU106364 ^64^), supporting the hypothesis that such unique OTUs in eggs may represent non-symbiotic, potentially opportunistic symbionts. Further work is needed to determine what ecological roles these symbionts potentially fulfil and their systematic relationships. For example, a high number of “uncultured” types suggest considerable taxonomic uncertainly, as has been observed for clade E *Symbiodinium* (see discussion in ^65^).

Maternal environmental effects, such as lipid contributions by dams, have well known effects on the early life stages of many marine organisms ^66^. However, our Bayesian models were not significantly improved by the addition of dam identity, suggesting that significant heritability estimates are attributable to genetic effects and not due to maternal environmental effects ^35^ or cytoplasmic inheritance ^67^. Whilst we can only speculate about the exact mechanisms that are being inherited by offspring, likely candidates include those involved in recognition and immunity pathways ^42^, with cell-surface proteins playing an important role in the selection of specific *Symbiodinium* strains by coral hosts ^68–70^. For example, these may include Tachylectin-2-like lectins, which have been implicated in the acquisition of A3 and a D-type in *A. tenuis* ^43,71,72^. Indeed, suppression or modification of the immune response has often been implicated in the formation of *Symbiodinium-cnidarian* partnerships ^42,73,74^. Although this has not yet been demonstrated in corals, human studies have shown that immune system characteristics underpin heritable components of the genome ^75^ and at least 151 heritable immunity traits have been characterized, including 22 cell-surface proteins ^76^.

Juvenile corals may be primed to take up specific *Symbiodinium* types through the transfer of genetic machinery that results in a by-product(s) that ensures juveniles are colonized by beneficial types and prevents colonization by unfavourable symbionts through competitive exclusion (e.g., maternal imprinting controlled by offspring loci ^67^). Such by products may be akin to amino acids, which have been shown to regulate the abundances of *Symbiodinium* populations ^77^. Sugars have also been found to influence bacterial communities in corals ^78^ and may have similar roles in regulating *Symbiodinium* communities. Trehalose, in particular, has been identified as an important chemical attractant between *Symbiodinium* and coral larvae and may help to regulate the early stages of symbiosis ^79^. Human studies also provide examples of sugars (both maternal and offspring derived) that make infant intestines less habitable for harmful bacteria, setting up conditions for preferential colonization by favourable bacteria ^80^. Bacterial diversity in cnidarian hosts can also be modulated through the production of antimicrobial peptides ^36^ and bacterial quorum sensing behaviour ^81^. Although neither of these mechanisms has been explored with respect to the regulation of *Symbiodinium* in corals, similar host/symbiont by-products may be influential in the regulation of *Symbiodinium* communities.

Heritability estimates based on parent-offspring regression and Bayesian MCMC methods were similar in *A. tenuis* but not in *M. digitata.* Differences between the estimates of these two methods for *M. digitata* may be due to the purely maternal basis of inheritance in this species, with the slope of parent-offspring regressions potentially more accurate for traits that are transmitted following sexual reproduction involving two parents. Alternatively, Bayesian MCMC methods, which do not rely on phenotypic information of parents, and instead only utilize information on relatedness among offspring and co-variances between them in the phenotypic trait being measured, may be more robust to a variety of different reproductive modes across organisms. Furthermore, outplanting juveniles to only one location may have introduced bias into the regression-based estimates, causing juveniles and adults from the OI location to appear more similar, potentially because they were exposed to similar environmental pools of symbionts, compared to juveniles from PCB parents.However, concordance between Bayesian (which do not rely on parental phenotypic information) and regression-based estimates suggests that this bias is negligible. Standard errors calculated in heritability studies are normally large ^5^ but Bayesian MCMC methods are robust, as they allow for estimation of heritability and statistical support of that estimate directly from posterior distributions. Therefore, although credibility intervals calculated were large, high densities of posterior distributions around our heritability estimates signify that these values are the most probable compared to values at lower posterior densities. This Bayesian method for determining uncertainty is robust, especially compared to frequentist methods where standard errors are approximate ^5^.

In conclusion, results presented here provide new insights into the role of host genetics and inheritance in governing *Symbiodinium* communities in corals. This information is important for determining the potential for host-symbiont partnerships to evolve. Variability in the symbiont community within and among families and evidence that variation is heritable, as supported by the moderate to high heritability estimates found, corroborate the likelihood that adaptive change is possible in this important symbiotic community. These results may also aid in the development of active reef restoration methods focused on assisted evolution of hosts and symbionts, in which targeted traits with moderate to high heritability increase the efficacy of breeding schemes. Adaptive change through heritable variation of symbionts is therefore another mechanism that corals may use to contend with current and future stressors, such as climate change.

## Materials and Methods

### Experimental breeding design and sample collection

For crossing experiments, gravid colonies of the horizontally-transmitting broadcast-spawning coral *Acropora tenuis* were collected in 2012 and 2013 from the northern (Princess Charlotte Bay (PCB): 13°46’44.544”S, 143°38’26.0154”E) and central Great Barrier Reef (GBR) (Orpheus Island: 18°39’49.62”S, 146°29’47.26’E).

In 2012, nine families of larvae were produced by crossing gametes from four corals (OI: A-B, PCB: C-D) on 2 December following published methods ^82^. The nine gamete crosses excluded self-crosses (Supplementary Table S1). Larvae were stocked at a density of 0.5 larvae per ml in one static culture vessel per family in a temperature-controlled room set at 27°C (ambient seawater temperature). Water was changed one day after fertilization and every two days thereafter with 1μM filtered seawater at ambient temperature. To induce settlement, 25 settlement surfaces (colour-coded glass slides) were added to each larval culture vessel six days post-fertilization, along with chips of ground and autoclaved crustose coralline algae (CCA, *Porolithon onkodes* collected from SE Pelorus: 18°33’34.87”S, 146°30’4.87”E). The number of settled juveniles was quantified for each family, and then placed randomly within and among the three slide racks sealed with gutter guard mesh. The racks were affixed to star pickets above the sediments in Little Pioneer Bay (18°36’06.2”S, 146°29’19.1”E) 11 days post fertilization. Slide racks were collected 29 days later (11 January 2013), after which natural infection by *Symbiodinium* was confirmed with light microscopy. Juveniles from each cross were sampled (n = 6 - 240 juveniles/family, depending on survival rates), fixed in 100% ethanol and stored at -20°C.

In 2013, 25 families were produced from gamete crosses among eight parental colonies: four from PCB and four from Orpheus Island (full details of colony collection, spawning, crossing and juvenile rearing in ^82^ (Supplementary Table S2). Larvae were raised in three replicate cultures per family. Settlement was induced by placing autoclaved chips of CCA onto settlement surfaces, which were either glass slides, calcium carbonate plugs or the bottom of the plastic culturing vessel. Settlement surfaces with attached juveniles were deployed randomly, 19 days post fertilization, at the same location in Little Pioneer Bay as in 2012, and collected 26 days later. Samples of juveniles (n = 1 - 194 juveniles per family) were preserved and stored as in 2012.

Thirty-two gravid colonies of the vertically-transmitting broadcast spawner *Montipora digitata* were collected from Hazard Bay (S18°38.069’, E146°29.781’) and Pioneer Bay (S18°36.625’, E146°29.430’) at Orpheus Island on the 30^th^ of March and 1^st^ of April 2015. Colonies were placed in constant-flow, 0.5 μM filtered seawater in outdoor raceways at Orpheus Island Research Station. Egg-sperm bundles were collected from a total of nine colonies on the 4^th^ and 5^th^ of April, separated with a 100 μm mesh and rinsed three times. Individual eggs and adult tissue samples were then placed in 100% ethanol and stored at -20°C until processing.

### Sequencing of *Symbiodinium* ITS-2 in egg, juvenile and adult coral samples

The number of juveniles of *A. tenuis* sequenced from each of the 9 crosses in 2012 ranged from 2 - 29 individuals (average ± SE: 11.3 ± 3) (Supplementary Table S1) and a single sample from each parental colony was sequenced concurrently. In 2013, 1 - 21 *A. tenuis* juveniles (average ± SE: 8.6 ± 1) were sequenced from each of the 20 families (of the original 25) that survived field deployment (Supplementary Table S2). The adult samples sequenced included three samples per colony from Orpheus parents (from the edges and center of each colony) and one sample per colony for Princess Charlotte Bay parents. For *M. digitata,* 5 – 12 eggs per dam were sequenced, along with one sample per maternal colony.

DNA was extracted from juveniles of *A.tenuis* in 2012 and 2013 with a SDS method ^82^ (additional description in Supplementary Methods). For *M. digitata,* single egg extractions used the same extraction buffers and bead beating steps as described in ^82^, although without the subsequent washes and precipitation steps because of the small tissue volumes of single eggs ^83^. Library preparation, sequencing and data analysis were performed separately for 2012 and 2013 samples of *A. tenuis* and *M. digitata,* as described in ^82^. Briefly, the USEARCH pipeline (v. 7) ^84^ and custom-built database of all *Symbiodinium-specific* NCBI sequences were used to classify reads ^85,86^, with blast hits above an E-value threshold of 0.001 removed, as they likely represented non-specific amplification of other closely-related species within the Dinoflagellata phylum (Supplementary Table S3). Cleaned reads were clustered with the default 97% identity and minimum cluster size of 2 (thus eliminating all singleton reads), after which all reads were globally aligned to 99% similarity with gaps counted as nucleotide differences.

*Symbiodinium* databases suggest that hundreds of subclades and types exist within *Symbiodinium* clades ^87,88^. These subclades and types likely represent distinct *Symbiodinium* species ^89^. However, the status of OTUs is less clear; they might represent either unique *Symbiodinium* genotypes or intragenomic variants ^10,17,18^ or both, but they are unlikely to represent distinct *Symbiodinium* species. Nevertheless, in some cases, OTUs map to known‘types’(see ^18^). Therefore, this OTU-based framework infers delineations between the OTU, subtype, and type levels ^18,89^. However, a large proportion of OTUs retrieved in this study are unlikely to represent intragenomic variants for two reasons. Firstly, the proportion of intragenomic variants retrieved as OTUs will depend on the methodology used to cluster sequence variants. Clustering across samples at 97% identity greatly diminishes retrieval of intragenomic variants ^18^. Secondly, in contrast to overestimating diversity, clustering across samples at 97% identity also results in an underestimation of relevant biological diversity ^90^. As there is no single-copy marker yet known for *Symbiodinium,* sequencing additional markers would result in intragenomic challenges similar to those found for ITS-2. Therefore, at this time, sequencing additional markers is not a panacea for dealing with intragenomic/multicopy variation. Finally, *Symbiodinium* OTUs listed as “uncultured” were assigned this term based on their Genbank NCBI identifiers, following verbatim the name given by the original depositors of these sequences. Quotes around the term were added to make clear that this is not a functional description or taxonomic designation. Analysis of rarefaction curves suggested that differences in sequencing depth across samples did not affect diversity estimates (additional description in Supplementary Methods).

### Data analysis and visualization

Sample metadata were mapped onto circular trait plots using the package ‘diverstree’ ^91^. To aid in visualizing the data on the *A. tenuis* plots, only OTUs that were found within at least three samples were kept, reducing the total OTU count from 422 to 134 for 2012 samples and from 568 to 181 for 2013 samples, giving an overall total of 315 OTUs for *A. tenuis.* To determine the overlap in *Symbiodinium* OTUs from *A. tenuis* data between years that were clustered and mapped separately, the 315 OTUs were aligned in Clustal OMEGA ^92^. OTUs that clustered and blasted to the same accession number (54 of the 315) were deemed to be the same OTU, resulting in a total of 261 distinct OTUs. In total, 80 unique OTUs were found in 2012, 127 were found in 2013, and 54 were shared between years.OTUs with a relative normalized abundance of less than 0.01% were classified as “background”, whilst those with abundances greater than 0.01% were considered “principal.” Rare, background types can play an important role in recovery post-bleaching caused by both low and high temperatures by becoming dominant symbionts ^50^. The cut-off of 0.01% chosen to designate background abundances here is commonly used in microbial, deep sequencing studies examining rare taxa ^93–95^, and has been found to fall within the detection limits of deep sequencing for *Symbiodinium* ^17^. Furthermore, 0.01% represents approximately 100-200 cells per square cm ^57^, a density of symbionts that has been recognised as ecologically relevant. For example, a survey of four coral species on the GBR revealed clade D populations existed, on average, at levels of 100-10,000 cells per cm^2 58^. This study is also the first to use deep sequencing to identify *Symbiodinium* communities in eggs and juveniles of corals, and therefore this lower threshold enabled the inclusion of a greater percentage of *Symbiodinium* communities with which to explore the diversity present in this life stage. OTUs were further classified by ubiquity across samples, whereby “core” OTUs were defined as those found in >75% of samples, “common” were found in 25 -75% of samples, and “rare” were found in < 25%. As far fewer OTUs were recovered from *M. digitata*samples, all 101 OTUs from the one year sampled were used to visualize and classify them by abundance and ubiquity, as described above. Differential abundance testing was performed with ‘DESeq2’, with Benjamini-Hochberg p-adjusted values at 0.05 ^96–98^.Networks and heatmaps were constructed using un-weighted Unifrac distances of the normalized *Symbiodinium* abundances in eggs only, where maximum distances were set at 0.4.

### Heritability analyses

We estimated the effects of host genotype and maternal environment on variation in *Symbiodinium* diversity using established quantitative genetic methods ^5,31^. The extent to which a trait (such as the host’s *Symbiodinium* community) is genetically regulated can be represented by the degree to which individuals share the same genes ^31^. The degree to which individuals share the same genes can be determined in at least two ways: 1) using information on relatedness through the construction of known pedigrees based on either reproductive crosses (as we have done here), twin data, or known breeding lines; or 2) using genomic marker data (Quantitative Trait Loci) ^30,99,101^. For example, whilst the full genome structure of twins is often not known in heritability studies, twin studies provide subjects of known relatedness (full sibs, half sibs), from which host-genotype sharing can be calculated. Therefore, we have constructed pedigrees of known relatedness using diallel and half-diallel cross designs to construct the degree to which individuals share the same genes (host genotype information). Pedigrees of known relatedness were then combined with host phenotypes for the trait *“Symbiodinium* community” that had been determined through sequencing. The *Symbiodinium* community is not a “proxy” for host-phenotype; it is the host phenotype for symbiosis. In a similar manner, host-phenotypes for symbiosis have been determined in studies of bacterial gut communities in insects and mammals ^39–41^. Heritability analysis therefore uses information on variation among samples in both host-genotype (calculated here through relatedness coefficients derived from pedigrees) and host-phenotype (i.e., *Symbiodinium* community determined through sequencing). We do not explicitly determine which elements of the host genotype regulate the variability in this host trait *(Symbiodinium* community); such a determination would require Quantitative Trait Loci analysis. Instead, our objective is to quantify the extent to which this trait is genetically regulated.

The *Symbiodinium* community associated with each adult, juvenile (A. *tenuis*) or egg *(M. digitata*) of the two coral species was characterized as a continuous quantitative trait of the host by converting community composition into a single diversity metric. Differences among juveniles in regards to their *Symbiodinium* communities were examined as a host phenotypic trait. Collapsing complex assemblage data into a single diversity value (local diversity measure)^102^ was necessary to apply a univariate heritability statistic. Such single diversity metrics have been used to explore the impact of host-genetic variation on bacterial symbiont populations residing within hosts across a range of environments in the adult and infant human body^39,41^ as well as in insects ^40^. The Leinster and Cobbold diversity metric (D) incorporates variance-normalized OTU abundances from linear models using negative binomial distributions, OTU sequence diversity, and OTU rarity in the following equation ^102^:

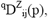

where “q” is a measure of the relative importance of rare species from 0 (very important) to ∞ (not important), and Z is a matrix of genetic similarities of OTUs i through j. Pairwise percent similarities between OTUs sequences were calculated in ‘Ape’ with a “raw” model of molecular evolution, in which the simple proportion of differing nucleotides between pairwise comparisons is calculated and no assumption is made regarding the probability of certain nucleotide changes over others. Finally, P is a matrix of normalized abundances corresponding to each sample and OTU. Incorporating both abundance and diversity of *Symbiodinium* types into heritability estimates is essential because changes in *Symbiodinium* community abundance dynamics can change the functional output of the symbiosis as a whole ^26^ and are important in determining coral resilience and bleaching susceptibility ^25,103,104^. Model inputs therefore take into account which OTUs were present or absent in each sample, OTU sequence diversity, and the abundance of each OTU.

Heritability estimates for both species presented here represent the initial *Symbiodinium* community with the time of sampling consistent with complete infection (i.e. defined by the presence of *Symbiodinium* throughout the polyp) of *A. tenuis* juveniles (19 - 22.5 days, personal observation, ^48,105^). Calculated heritability may vary among traits and throughout ontogeny (i.e. with body size ^106^) and hence we therefore make no predictions about the heritability of *Symbiodinium* communities at later ontogenic stages. However, as the early *Symbiodinium* community can influence juvenile survival ^82^and because we do not yet know how the earliest communities impact later ones, evaluating the heritability at this initial stage is a logical first step.

Two methods were used to assess heritability. Bayesian methods are powerful tools for assessing heritability of natural (i.e. non-lab, non-model) populations and for non Gaussian traits (see ^5^ for a full discussion of the advantages of using Bayesian inference in quantitative genetics). However, parent-offspring regressions were also calculated to facilitate comparisons with previous studies as they make up a majority of estimates available in the literature. The correspondence in heritability estimates between these two methods is well-established (e.g. h^2^=0.51 vs 0.52 for *Drosophila melanogaster* traits ^31,107^), although Bayesian MCMC estimates are generally lower ^5^ and confidence intervals around mean estimates generally smaller, especially at low levels of heritability ^108^. Importantly, neither method is dependent on the known relatedness of the parents, but instead rely on relatedness among the juveniles themselves (sib analysis comprised of full and half sibs) or comparisons between juveniles and adult phenotypes (parent-offspring regressions) ^30^.

#### Regression-based estimates of heritability

Phenotypic values of offspring can be regressed against parental midpoint (average) phenotypic values, with the slope being equal to the narrow-sense heritability of the trait of interest ^31,32^. Parental midpoint values were calculated by taking the average of dam and sire *Symbiodinium* diversities for each family and then regressing these values against diversity values for the offspring of each family. Precision of the heritability estimate increases when parents vary substantially in the trait of interest ^31^. Coral colonies dominated by a single or mixed *Symbiodinium* communities (C, D, C/D communities) can be considered biological extremes and ample evidence describes their contrasting physiological impacts on coral hosts (i.e., growth, bleaching) when associated with D versus C communities in particular ^26^. Therefore, parental colonies selected for breeding were dominated by C1 (families W5, 10) or had mixed communities of C1/D1 (W7), C1/D1/D1a (W11, PCB4, 6, 8, 9), or multiple A, C1 and D types (OI3, 4, 5, 6) (Fig. 2b).

#### Bayesian linear mixed model estimates of heritability

Heritability estimates were derived from estimates of additive genetic variance calculated from the ‘animal model,’ a type of quantitative genetic mixed effects model incorporating fixed and random effects, and relatedness coefficients amongst individuals ^109^. The animal model was implemented using Bayesian statistics with the package ‘MCMCglmm’ ^110^. The model incorporated the diversity metric calculated for each juvenile and the pedigree coefficient of relatedness as random effects. Bayesian heritability models were run with 1.5 x10^6^ iterations, a thinning level of 800 (*A. tenuis*) or 250 *(M. digitata),* and a burn-in of 10% of the total iterations. A non informative flat prior specification was used, following an inverse gamma distribution ^35^. Assumptions of chain mixing, normality of posterior distributions and autocorrelation were met. The posterior heritability was calculated by dividing the model variance attributed to relatedness by the sum of additive and residual variance. The impact of environmental covariance **(VEC)** was reduced by randomly placing families within the outplant area ^31^. Maternal environmental effects were assessed and were not significant for either *A. tenuis* or *M. digitata* based on Deviance Information Criteria (DIC) from Bayesian models ^35^. The influence of different settlement surfaces for *A. tenuis* juveniles in 2013 was assessed using linear mixed models (fixed effect: substrate, random effect: family) in the ‘nlme’ package ^111^ using the first principal component extracted from PCoA plots and incorporating weighted Unifrac distances of normalized *Symbiodinium* abundances for juveniles. Model assumptions of homogeneity of variance, normality, and linearity were met. Substrate type did not significantly explain *Symbiodinium* community differences among samples (LME: F(_4_)= 1.05, *p =* 0.38).

#### Impact of intragenomic variation on heritability analysis

The multicopy nature of *Symbiodinium* genomes and the presence of intragenomic variants make taxonomic assignments for distinct *Symbiodinium* sequences difficult, however, advances have been made to name and elucidate the functional diversity within *Symbiodinium* ^112–115^. Single base pair variations in key genetic regions (e.g., intragenomic spacer region-2 ITS-2) can be the sole difference between important taxonomic entities, for example, between a new thermally tolerant C3 type (S. *thermophilum*) and the ubiquitous C3 type ^116^; which further highlights the need for sensitive methodologies. Whilst different methods have been used to incorporate intragenomic variation into *Symbiodinium* taxonomy designations (i.e. single cell sequencing ^117^, and pairwise correlations ^10,17,18^), the combination of single-cell sequencing, gel-based methods and next generation sequencing suggest that clustering at 97 % sequence similarity (the cut-off used here), is sufficient to collapse *Symbiodinium* from clades A, B and C into type-level designations ^18^.

Even without accounting for intragenomic variation using the 97% clustering threshold, heritability analysis should be impacted little by these pseudo-variants given that intragenomic variants are found within the same genome. These groups of variants would therefore be inherited together and do little to impact variance between individuals of different families (which are important for calculating heritability), causing the bias in a systematic manner. To test this, we employed a three-step approach previously used to classify intragenomic variants ^82^ to the *M. digitata* dataset. Initial groups of OTUs were chosen from those that clustered closely together on the dendrogram as they have higher per cent similarity relative to other sequences. Correlation coefficients for these groups of closely clustered OTUs were then calculated, and OTUs having highly positive or negative correlations coefficients (-1 to -0.8, 0.8 to 1) were identified as candidate intragenomic variants. To test the impact of accounting for intragenomic variants on Bayesian heritability analysis, MCMC models were then re-run the same way as described above but now incorporating intragenomic variants into the new-derived diversity metric.

## Acknowledgements

We would like to thank Margaux Hein, Mikhail Matz, Marie Strader, Greg Torda, Sarah Davies, Natalie Andrade, Tess Hill and the staff at Orpheus Island Research Station for help with field work and spawning at Orpheus Island. We also thank Dr. Ray Berkelmans and the crew on the RV Ferguson for help with the collection of corals from the northern GBR.

All samples of *A. tenuis* from Wilkie Island and Orpheus Island and *M. digitata* from Orpheus Island were collected under Great Barrier Reef Marine Park Authority permits: G12/35236.1, G13/36318.1, and G10/33312.1. Funding was provided by the Australian Research Council through ARC CE1401000020, ARC DP130101421 to B.L.W. and AIMS to L.K.B.

## Author Contributions

K.M.Q., B.L.W., and L.K.B. designed and conducted the experiments, K.M.Q. analysed the data and wrote the manuscript, and all authors made comments on the manuscript.

## Data availability statement

All raw sequencing data will be deposited in the NCBI Sequence Read Archive under Accession number SRX.

